# Reversible auto-inhibitory regulation of *Escherichia coli* metallopeptidase BepA for selective β-barrel protein degradation

**DOI:** 10.1101/2020.07.07.192476

**Authors:** Yasushi Daimon, Shin-ichiro Narita, Ryoji Miyazaki, Yohei Hizukuri, Hiroyuki Mori, Yoshiki Tanaka, Tomoya Tsukazaki, Yoshinori Akiyama

## Abstract

*Escherichia coli* periplasmic zinc-metallopeptidase BepA normally functions by promoting maturation of LptD, a β-barrel outer membrane protein involved in biogenesis of lipopolysaccharides, but degrades it when its membrane assembly is hampered. These processes should be properly regulated to ensure normal biogenesis of LptD, but the underlying mechanism of regulation, however, remains to be elucidated. A recently solved BepA structure has revealed unique features, in particular the active site is buried in the protease domain and conceivably inaccessible for substrate degradation. Additionally, the His-246 residue in the loop region containing helix α9 (α9/H246 loop), which has a potential flexibility and covers the active site, coordinates the zinc ion as the fourth ligand to exclude a catalytic water molecule, thereby suggesting that the crystal structure of BepA represents a latent form. To examine the roles of the α9/H246 loop in the regulation of the BepA activity, we constructed BepA mutants with a His-246 mutation or a deletion of the α9/H246 loop and analyzed their activities *in vivo* and *in vitro*. These mutants exhibited an elevated protease activity and, unlike the wild-type BepA, degraded LptD that is in the normal assembly pathway. In contrast, tethering of the α9/H246 loop repressed the LptD degradation, which suggests that the flexibility of this loop is important to the exhibition of the protease activity. Based on these results, we propose that the α9/H246 loop undergoes a reversible structural change that enables His-246-mediated switching (histidine switch) of its protease activity, which is important for regulated degradation of stalled/misassembled LptD.

## Introduction

Living organisms defend themselves from the hazardous environmental stresses in various ways. In gram-negative bacteria, the outer membrane (OM) acts as a barrier to toxic compounds including chaotropic reagents, detergents and antibiotics (1). The OM is a lipid bi-layer membrane as other biological membranes, but is unique in that its outer leaflet is composed of lipopolysaccharides (LPSs) that are important to maintain OM integrity and confer a cell resistance to toxic compounds (1, 2). LPSs are synthesized and maturated on the cytoplasmic side of the inner membrane, flipped across the inner membrane by MsbA, and transported to the OM by the Lpt proteins (2). At the final step of the LPS transport, a complex of β-barrel OM protein (OMP) LptD and lipoprotein LptE catalyzes insertion of LPS into the OM as the LPS translocon (2, 3). The LptD/E hetero-dimer has a peculiar structure wherein the protein domain of LptE is accommodated within the membrane-embedded β-barrel domain of LptD (4–6). OMPs are also crucial to maintaining the OM structure and functions. Defective biogenesis of β-barrel OMPs causes increased drug sensitivity of a cell. They are synthesized as a precursor with a cleavable signal peptide and translocated to the periplasm by the Sec translocon. OMPs are then exported to the OM by the aid of periplasmic chaperones/foldases (7, 8) and inserted into the OM by the function of the β-barrel assembly machinery (BAM) complex, the OMP translocon in the OM. The BAM complex is composed of 5 components; one β-barrel OMP (BamA) is associated with 4 lipoproteins (BamB-E) (8, 9). This complex has a silk-hat (top-hat) like structure wherein the β-barrel domain of BamA corresponds to the OM-embedded “crown” and the periplasmic “brim”, which is supposed to undergo dynamic structural changes, is formed by the periplasmic POTRA domains of BamA and the lipoprotein components (9–11).

*Escherichia coli* BepA, a periplasmic M48 family metallopeptidase belonging to gluzincins (12), is required to maintain the functional integrity of the OM possibly through promotion of biogenesis and quality control of LptD (13–16). LptD has four Cys residues and its mature form (LptD^NC^) contains two disulfide bonds that are formed by non-consecutive pairs of the Cys residues (C31-C724 and C173-C725) (17). However, it has been shown that just after transport to the periplasm, an assembly intermediate (LptD^C^) with consecutive disulfide bonds (C31-C173 and C724-C725) was first formed and converted to LptD^NC^ (13, 18). This conversion is accelerated by BepA. In addition, when a normal assembly of LptD is blocked by a mutation in *lptD* (*lptD4213*) (19), or decreased availability of LptE, BepA degrades the stalled or misassembled LptD (13, 16). BepA is composed of two domains (13–15) an N-terminal M48 family metallopeptidases domain (12) and the C-terminal tetratricopeptide repeat (TPR) domain (20–22). BepA has been shown to interact with the BAM complex components and LptD through its TPR domain (13, 14). These interactions are important for the functions of BepA in the biogenesis and quality control of LptD (14). Furthermore, previous studies of the BepA TPR domain suggested that this domain is inserted into the interior of the periplasmic ring-like structure (“brim”) of the BAM complex (14, 15).

Recently, we determined the structure of the full-length BepA (Fig. 1*A*, PDB ID: 6AIT) (15). The overall structure of BepA coincided well with the structure of the minigluzincin that represents a minimal structural scaffold of the catalytic domains of the M48 and M56 family metallopeptidases (23), except that it has several unique regions including those containing α6 and α9 helices (Fig. S1). The α6 and α9-containing loop regions cover the protease active site, apparently hindering access of substrates to the active site. Gluzincins have an active site zinc ion that is generally coordinated in a tetrahedral way by three ligand residues (two His in the HExxH motif and the third ligand (typically Glu)) and a solvent water molecule required for the catalysis (24). BepA has two His residues (His-136 and His-140) in the conserved HExxH motif in the α4 helix and Glu-201 in the α7 helix that serve as ligands for zinc coordination. In gluzincins, the catalytic water is generally bound to the zinc ion and Glu in the HExxH motif (Glu-137 in BepA) that acts as the general base/acid (24). Intriguingly, the BepA structure showed that it does not have the catalytic water molecule, but instead has the fourth zinc ligand residue (His-246 in the α9-containing loop) (Fig. 1*B*). The solved structure suggests that BepA in this state is a latent form with its protease activity repressed, which is consistent with the very low *in vitro* proteolytic activity of purified BepA (13). To understand the functions of BepA, it will be important to know how His-246 is involved in the exhibition of the protease activity of BepA.

**Fig. 1.**
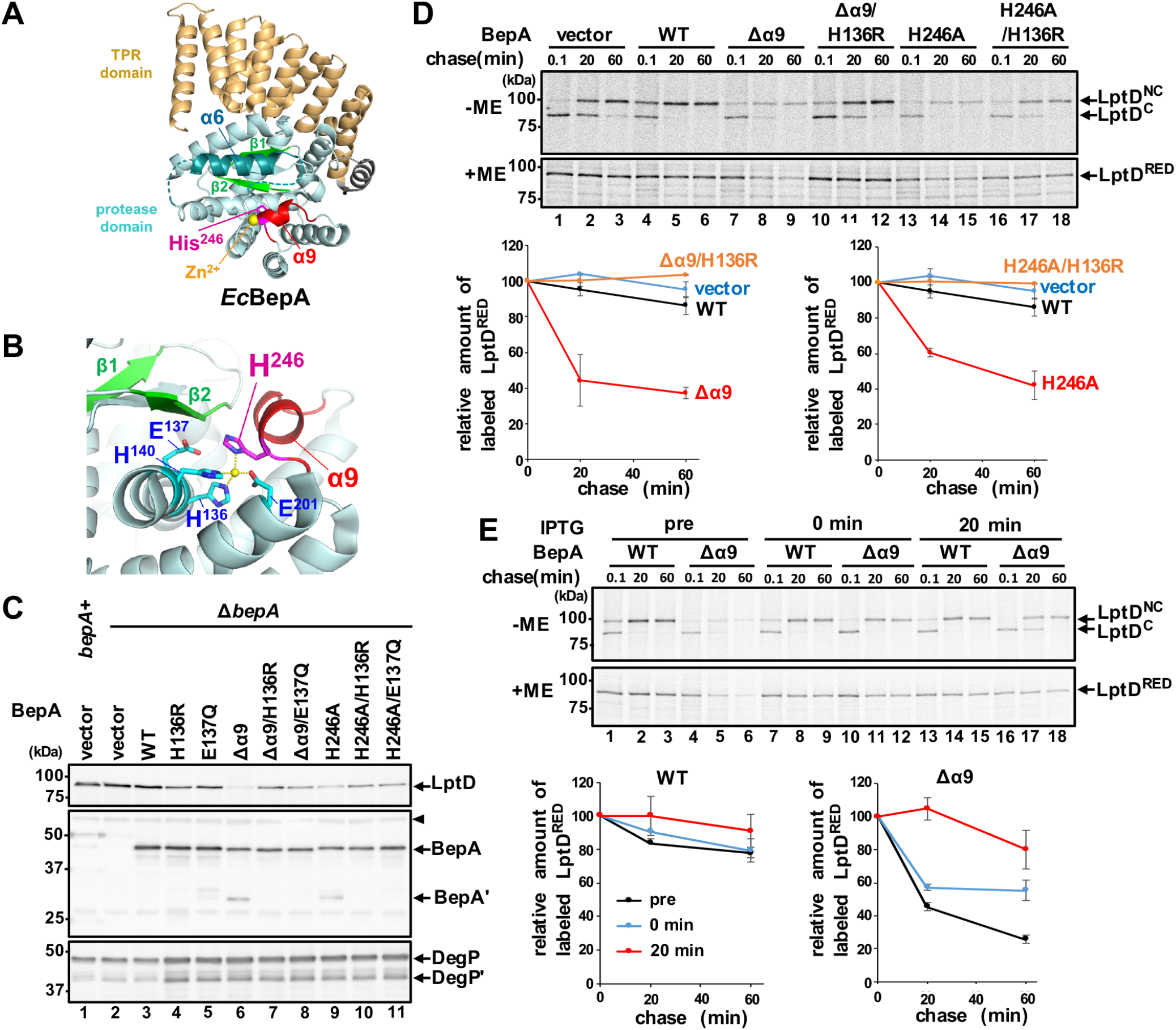
Degradation of chromosomally-encoded LptD by BepA(Δα9) and BepA(H246A). (*A*) The crystal structure of BepA (PBD ID: 6AIT). The α9/H246 loop (red), α6 loop (teal), β1/β2 (light green), zinc atom (yellow) and His-246 residue (magenta) are indicated. The TPR domain is shown in orange. Broken lines indicate disordered regions. (*B*) A close view of the proteolytic active site of BepA. The catalytic residues are shown in cyan. (*C*) Accumulation levels of chromosomally-encoded LptD in cells expressing the Δα9/H246 loop mutants. Wild-type cell carrying pUC18 (*bepA*^+^) and Δ*bepA* cells carrying either pUC18 (vector), pUC-bepA (WT), or its derivatives were grown at 37°C in M9-based medium with 1 mM IPTG. Total cellular proteins were analyzed by immunoblotting with anti-LptD (upper panel), anti-BepA (middle panel), or anti-DegP (lower panel) antiserum. BepA' and DegP' indicate a degradation product of BepA and DegP, respectively. The arrowhead indicates a non-specific band serving as a loading control. The migration positions of molecular weight markers are shown. (*D*) Pulse-chase analysis of LptD. Δ*bepA* cells carrying either pUC18 (vector), pUC-bepA, or its derivatives were grown at 37°C in M9-based medium, induced with 1 mM PTG for 15 min, pulse-labeled with [^35^S]-Met for 3 min and chased with excess unlabeled Met for the indicated times. Proteins were immunoprecipitated with anti-LptD antiserum and analyzed by nonreducing (−ME) or reducing (+ME) SDS/PAGE. Relative band intensities of total LptD (LptD^RED^) are shown in the lower graphs (the values for the band intensity at 0.1 min chase was set to 100), in which mean values of two independent experiments are shown with standard deviations. (*E*) Effects of BepA(Δα9) expression on the stability of mature LptD (LptD^NC^). Pulse-chase analysis of LptD were conducted as in *D* except that expression of BepA or BepA(Δα9) were induced at 15 min before pulse-labeling (pre), simultaneously with the initiation of chase (0 min), or at 20 min after the initiation of chase (20 min). Relative band intensities of total LptD are depicted as in *D*.

In this study, we examined the roles of His-246 and the loop region containing α9 and His-246 in the regulation of the BepA protease activity expression. Our results suggest that the proteolytic activity of BepA is usually repressed by His-246 but expressed as a result of the movement of the α9 loop region that will dislocate His-246 from the catalytic zinc ion. We propose that the proteolytic activity of BepA is reversibly regulated through binding of His-246 to the zinc ion, which is important for the selective degradation of misassembled or stalled LptD.

## Results

### The His-246 residue of BepA is required for the regulated expression of the proteolytic activity

Our previous X-ray structural study of BepA showed that in addition to the three zinc-coordinating residues (two His in the H^136^ExxH motif and Glu-201) conserved among gluzincins, His-246 located adjacent to the C-terminus of a short helix α9 (Ile-242 to Leu-244) coordinates the active site zinc ion in place of a zinc bound water molecule that is required for catalysis (15) (Fig. 1*A* and *B*). This suggests that the protease activity of BepA is auto-inhibited and that some conformation changes that displace His-246 will be needed for BepA to exhibit its protease activity in a timely manner.

Examination of the crystal structure of the a *Geobacter sulfurreducens* BepA homolog (Q74D82; PDB ID 3C37) showed that it has a very similar architecture to the protease domain of *E. coli* BepA, with His-208 corresponding to His-246 in *E. coli* BepA coordinating zinc ion being quite similar to *E. coli* BepA, except that a region including α9 is disordered (Fig. S2). It would thus be conceivable that the loop region containing α9 and H246 (designated as α9/H246 loop hereafter) of BepA (shown in red in Fig. 1*A* and *B*) potentially has a dynamic nature. Also, comparison of the secondary structure arrangement of the BepA protease domain with that of the proposed minimal scaffold of the minigluzincin (23) revealed that the latter lacks the structural element corresponding to α9 of BepA (Fig. S1), thereby suggesting that α9 is not directly required for the protease activity of BepA. Based on these considerations, we hypothesized that the α9/H246 loop has structural flexibility, which allows His-246 to act as an ON/OFF switch for regulation of the protease activity of BepA.

We reasoned that, if His-246 is indeed involved in repression of the BepA protease activity through coordination of the zinc ion, a mutational alteration of His-246 or removal of the α9/H246 loop would activate the protease function of BepA. We thus constructed a mutant form of BepA either having an H246A mutation or a deletion of the α9/H246 loop (Δα9; the amino acid residues from Pro-239 to Pro-247 were replaced by a flexible linker sequence Gly-Ser-Gly-Ser-Gly-Ser) and examined their effects on the accumulation level of the chromosomally-encoded LptD protein in a Δ*bepA* strain (Fig. 1*C*). We found that expression of the Δα9 or the H246A mutant greatly reduced the accumulation level of LptD, although, as shown previously (14), expression of wild type BepA little affected it. The results also showed that introducing an additional mutation in the protease active site motif (H136R or E137Q) that compromises the proteolytic activity of BepA largely canceled the effects of the Δα9 or H246A mutation. This was evidenced by the increased accumulation of LptD by the strains expressing the protease-dead derivatives of the Δα9 or H246A mutant as compared with the same mutants with the intact active site. The Δα9 and H246A derivatives of BepA accumulated at slightly lower levels than the control proteins without these mutations. Notably, in addition to the full length proteins, a fragment (~30 kDa) was detected with anti-BepA antibodies for BepA(Δα9) and BepA(H246A). This fragment should be produced by self-cleavage because no such fragment was observed for their derivatives carrying the H136R or E137Q mutation.

The above results suggest that BepA(Δα9) and BepA(H246A), but not wild type BepA, can degrade LptD even in a normal strain (i.e. in the absence of LptD assembly failure). We noticed that expression of the mutant forms of BepA carrying the Δα9 or H246A mutation and/or the active site mutations caused an elevated accumulation of DegP, suggesting that it induces extracytoplasmic stress responses (25). However, the observed decreases in the levels of LptD upon expression of BepA(Δα9) and BepA(H246A) should not result from the induction of the extracytoplasmic stress responses, as the similar levels of DegP were accumulated upon induction of BepA(Δα9) and BepA(H246A), and their protease-dead derivatives (Fig. 1*C*). We also occasionally observed that the accumulation levels of LptD decreased substantially upon expression of the E137Q mutant, presumably due to the lowered expression of LptD caused by some unknown reasons. The H136R mutants were, therefore, used as the protease-dead BepA derivatives in the following experiments. To minimize the possible indirect effects due to the BepA mutant overexpression and examine BepA-dependent LptD degradation in more detail, we next examined the maturation and stability of chromosomally-encoded LptD by pulse-chase experiments after short (15 min) induction of BepA and its derivatives (Fig. 1*D*). Cells expressing each protein were pulse-labeled with [^35^S]-Met for 3 min and chased with unlabeled Met for 0.1, 20, or 40 min at 37°C. LptD was then recovered by immunoprecipitation. Because LptD is converted during the assembly process from the intermediate form (LptD^C^). having consecutive disulfide bonds. to the mature form (LptD^NC^). having non-consecutive ones (13, 18), a portion of the samples were analyzed by non-reducing SDS/PAGE to monitor the maturation process (LptD^C^ is electrophoresed faster than LptD^NC^ in a non-reducing condition). Parallel non-labeling experiments conducted under essentially the same condition showed similar levels of accumulation for BepA and its derivatives as well as a milder DegP induction (Fig. S3*A*). The results with the reducing SDS/PAGE (+ME) (Fig. 1*D*, lower panel) showed that LptD was destabilized significantly by expression of BepA(Δα9) or BepA(H246A), whereas it was stable over 40 min when wild type BepA was expressed. This destabilization depended on the intactness of the protease active sites as LptD remained stable upon expression of BepA(Δα9/H136R) and BepA(H246A/H136R), thereby confirming the results of the immunoblotting analyses (Fig. 1*C*). Analysis with the non-reducing SDS/PAGE (-ME) (Fig. 1*D*, upper panel) showed that a small amount of LptD^NC^ that was generated in the presence of BepA(Δα9) or BepA(H246A) remained stable during the chase periods, suggesting that LptD^NC^ is not susceptible to the Δα9 or H246A mutant. To further examine this, BepA(Δα9) was induced at the same time as cold Met addition or 20 min after the initiation of chase (Fig. 1*E*). The results showed that the induction of not only the wild type BepA, but also BepA(Δα9) during the chase had very little effect on the stability of LptD^NC^. Similar amounts of BepA accumulated after 15 min induction under these conditions (Fig. S3*B*). These results collectively suggest that, while LptD in the normal biogenesis pathway is not susceptible to degradation by wild type BepA, BepA(Δα9) and BepA(H246A) can degrade an intermediate (most likely LptD^C^) at an early phase of its OM assembly, although they can not degrade OM-assembled LptD with the mature conformation (LptD^NC^).

### BepA with the deletion of the α9/H246 loop or the H246A mutation exhibits increased proteolytic activity in vitro

To examine the protease activity of BepA(Δα9) and BepA(H246A) directly, these proteins were purified by affinity chromatography using a C-terminally attached His_10_-tag (Fig. 2*A*). We found that BepA(Δα9) and BepA(H246A) underwent considerable self-degradation during purification and that introduction of A181E/L182T mutations (in the region just following α6) greatly suppressed the self-degradation (Fig. 2*A* and *Supplementary Results*). The purified preparations of BepA(Δα9) and BepA(H246A) carrying the A181E/L182T mutations (represented as BepA(Δα9)* and BepA(H246A)*, respectively) contained primarily full-length proteins (Fig. 2*A*, lanes 7 and 13), although incubation of these proteins for 8 h at 37°C still resulted in decrease in the amount of the full-length proteins and concomitant generation of faster migrating species. This conversion should also result from self-degradation because it was not observed with the BepA(Δα9)* and BepA(H246A)* derivatives additionally having the active site mutation (H136R) (Fig. 2*A*), or when incubation was performed in the presence of a metal chelating reagent such as 1,10-phenanthroline or EDTA, inhibitors of zinc metallopeptidases (Fig. S4*A*). N-terminal sequence analysis of a major self-degradation product of BepA(Δα9)* (Fig. 2*A*, green arrowhead) indicated that it was cleaved between Ala-171 and Met-172. We used these preparations in addition to the “wild type” BepA carrying the A181E/L182T mutations (BepA*) as well as their protease-dead (H136R) derivatives for *in vitro* degradation assay with a model substrate α-casein described below.

**Fig. 2.**
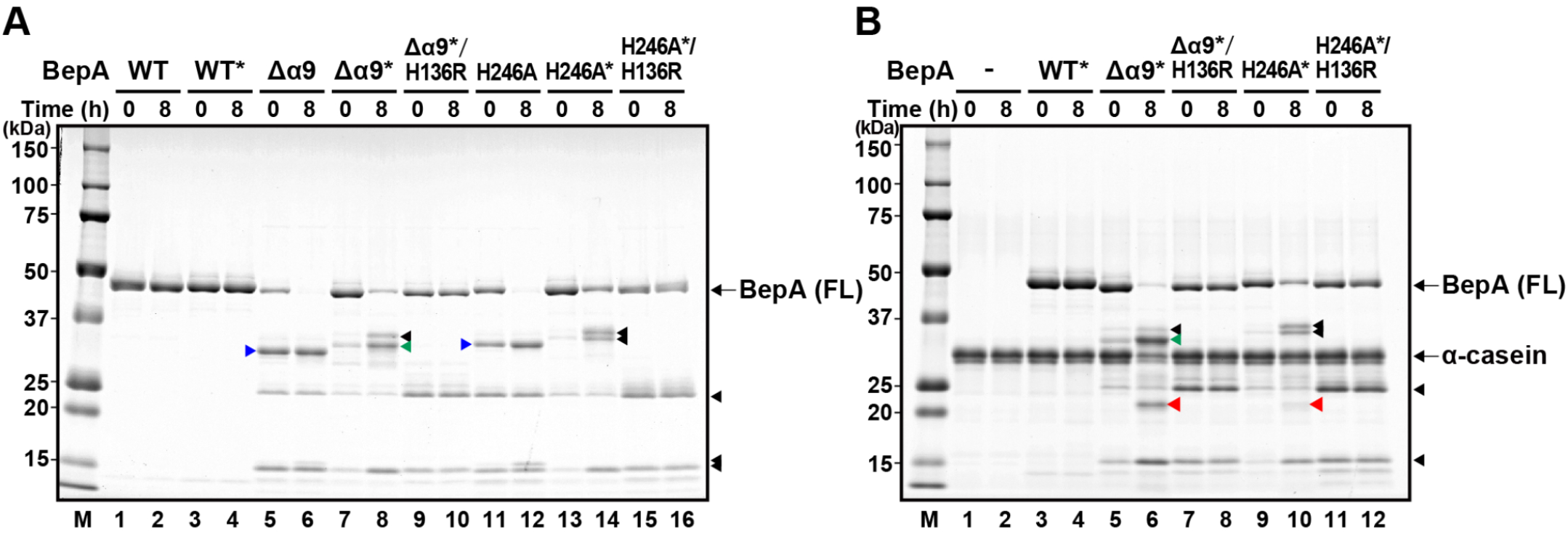
BepA with the deletion of the α9/H246 loop or the H246A mutation exhibits increased proteolytic activity *in vitro*. (*A*) Self-degradation of the Δα9 and the H246A mutants. Wild-type (WT) or the indicated mutant forms of BepA with a C-terminal His10-tag were incubated at 37°C for 0 or 8 h and analyzed by SDS/PAGE and CBB staining. BepA derivatives with the A181E/L182T mutations are indicated by asterisks. Full-length (FL) BepA derivatives are indicated. (*B*) Degradation of α-casein by the BepA derivatives. Wild-type (WT) or the indicated mutant forms of BepA, carrying the A181E/L182T mutations (indicated by asterisks) and a C-terminal His10-tag, were incubated with α-casein at 37°C for 0 or 8 h and analyzed by SDS/PAGE and Coomassie Brilliant Blue G-250 (CBB) staining. Full-length (FL) BepA derivatives and α-casein are indicated. Proteolytic fragments of α-casein are indicated by red arrowheads. The C-terminal fragments of BepA derivatives generated by cleavage between A181 and L182 and between A171 and M172 are indicated by blue and green arrowheads, respectively. Other proteolytic fragments of BepA derivatives are indicated by black arrowheads. The migration positions of molecular weight markers are shown on the left.

We previously reported that BepA showed a very weak proteolytic activity against α-casein, yielding a small amount of a cleaved fragment only after a long (24 h) incubation with the purified wild-type BepA (13). The α-casein cleavage was not observed after 8 h (Fig. 2*B*) or 24 h (Fig. S4*B*) incubation with the wild-type BepA prepared in this study. The exact reason for the failure in detection of the caseinolytic activity with the purified BepA this time remains unclear, but this might be ascribed to the very low activity of BepA that is just around the threshold of detection and greatly affected by subtle differences in the purified protein preparations. In marked contrast to wild type BepA, 8-h incubation of α-casein with the BepA(Δα9)* or BepA(H246A)*, which were purified parallelly with wild type BepA*, substantially decreased the amount of full-length α-casein (Fig. 2*B*) with concomitant generation of a α-casein fragment of ~22 kDa (red arrowheads). The effect on α-casein was more marked with BepA(Δα9)* than with BepA(H246A)*. It should have resulted from degradation of α-casein by these BepA mutants as the ~22 kDa fragment was not observed with their derivatives additionally having the protease-dead mutation (H136R) (Fig. 2*B*), or in the presence of a metal chelating reagent, 1,10-phenanthroline or EDTA, inhibitors of metallopeptidases (Fig. S4*C*). Comparison of the α-casein degradation by BepA(Δα9) and BepA(H246A) with or without the A181E/L182T mutations showed that the A181E/L182T mutations had little effect on the proteolytic activity of the Δα9 and the H246A mutants (Fig. S4*D*). The *in vitro* results with the purified proteins clearly demonstrated that BepA(Δα9)* and the BepA(H246A)* exhibit elevated protease activity as compared with wild type BepA*. The *in vivo* and *in vitro* results collectively suggest that the α9/H246 loop acts to repress the protease activity of BepA, and once this repression was released, BepA degrades normally assembling LptD.

### Tethering of the helix α9 inhibits the protease activity of BepA

BepA lacking the α9/H246 loop or with the H246A mutation degraded LptD that was assembling through a normal biogenesis pathway *in vivo* (Fig. 1) and exhibited an elevated protease activity *in vitro* (Fig. 2). As we hypothesized, the mobility of the α9/H246 loop could be important for regulation of the BepA protease activity via His-246. To investigate this hypoesthesia, we examined whether immobilization of the the α9/H246 loop by disulfide cross-linking affects the protease activity of BepA. In the structure of full-length BepA (15), Glu-241 located adjacent to the N-terminus of α9 is close (about 3.6 Å) to Glu-103 in the loop between the β1 and β2 strands (Fig. 3*A*). These two residues are individually or simultaneously replaced by a Cys residue. To examine the formation of the intramolecular disulfide bond between the introduced Cys residues and the protease activity of these mutants, we used the LptD overproducing system (14). In this system, most of the overexpressed LptD molecules failed to associate with the chromosomally-encoded LptE as LptE became limiting, and as a result, they are degraded by co-expressed BepA depending on the intactness of the active site to yield degradation products (Fig. 3*B*, lane 2 and Fig. 3*C*, lane 2). Co-expression of each of the single Cys mutants resulted in essentially the same results with wild type BepA, whereby reduced accumulation of full-length LptD and generation of the degradation products were observed (Fig. 3*B*, lanes 4 and 5). In contrast, LptD degradation was hardly observed with the double Cys (E103C/E241C) mutant (Fig, 3*B*, lane 6; 3*C*, lane 4). The E103C/E241C mutant migrated faster in non-reducing SDS/PAGE (-ME) as compared with in reducing SDS/PAGE (+ME), indicating that the Cys residues in this mutant were mostly oxidized to form a disulfide bond. When cells expressing the E103C/E241C mutant were cultured in the presence of 1 mM Tris(2-carboxyethyl)phosphine (TCEP), a reducing reagent, the disulfide bond of BepA was mostly reduced and the LptD degradation products were generated (Fig. 3*C*, lane 8) albeit at lower levels as compared with cells expressing the wild type BepA (lane 6). This indicates that the inability of the oxidized E103C/E241C mutant to degrade LptD was at least partly brought about by the disulfide crosslinking between the Cys residues introduced at positions 103 and 241. Analysis of LptD with non-reducing SDS/PAGE (-ME) showed that under the condition used, mature LptD (LptD^NC^) was mostly resistant to reduction by TCEP, possibly because LptD^NC^ is associated with the OM with a stable conformation, although some of the assembly intermediate (LptD^C^) was converted to the reduced form (LptD^RED^). Wild type BepA and the E103C/E241C mutant degraded LptD^RED^ and LptD^C^, but not LptD^NC^, to produce the degradation products in the presence of TCEP (Fig. 3*C*). These results showed that the disulfide-bond mediated immobilization of the α9/H246 loop strongly hinders the protease activity of BepA.

**Fig. 3.**
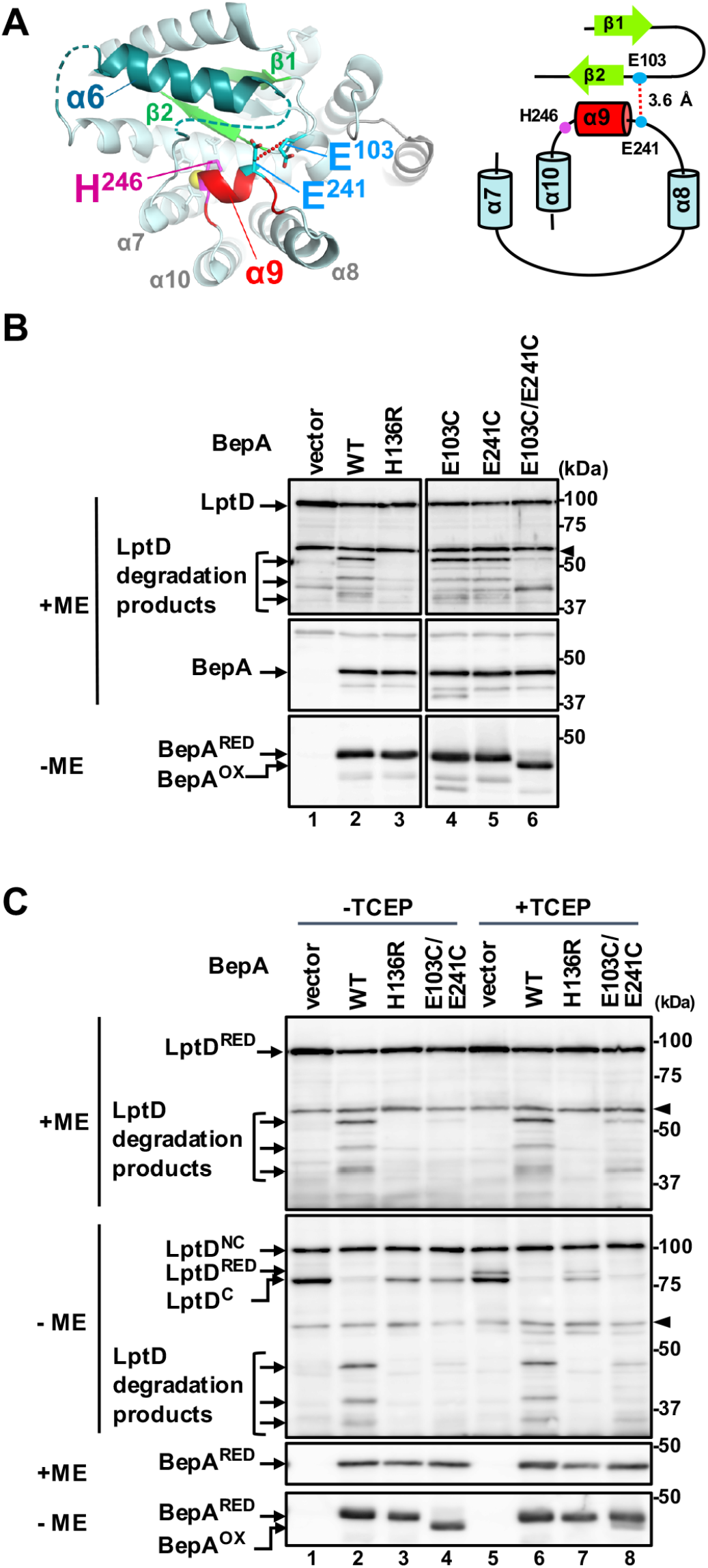
Tethering of the helix α9 inhibits degradation of overproduced LptD by BepA. (*A*) A closed view and a schematic representation of the regions connected by the disulfide bond formation between E103C and E241C (shown in cyan). The distance between β-carbons of Glu-103 and Glu-241 is shown. (*B*) LptD degradation by the α9/H246 loop-fixed mutants of BepA. Δ*bepA*/pTWV-lptD-his10 cells carrying an empty vector pSTD689 (vector) or either of the plasmids encoding the indicated BepA derivatives were grown in M9-based medium with 1 mM IPTG, and total cellular proteins were analyzed by reducing (+ME) or non-reducing (-ME) SDS/PAGE and immunoblotting with anti-LptD (upper panel) or anti-BepA (middle and lower panels) antiserum. A ~40 kDa band reacted with LptD antiserum in the lanes for the vector (lane 1) and E103C/E241C (lane 6) samples are unknown backgrounds that were detected occasionally. The arrowheads indicate non-specific bands serving as a loading control. (*C*) Effects of disulfide cleavage by TCEP on the LptD degradation by the E103C/E241C mutant. Cells were grown and analyzed as in *B* except that they were treated with 1 mM (final conc.) of TCEP for 10 min at an early log phase before sampling. The representative results of two independent replicates are shown. BepA^OX^ and BepA^RED^ indicate the disulfide-oxidized and reduced form of BepA(E103C/E241C), respectively.

### The α9/H246 loop is not essential for the chaperone-like activity of BepA

BepA has a chaperone-like activity that promotes maturation of LptD, as revealed by accelerated conversion of LptD^C^ to LptD^NC^ (13, 14). Pulse-chase experiments (Fig. 1*D* and *E*) showed that the conversion appeared to be slightly faster when wild type BepA was co-expressed as compared with BepA(Δα9) and BepA(H246A). However, self-cleavage of the Δα9 and the H246A mutants, which would affect the accumulation levels of these proteins, as well as unregulated degradation of LptD by them, makes the evaluation of their chaperone-like activity complicated. We thus examined the chaperone-like activity by pulse-chase experiments using the protease-dead derivatives of BepA, BepA(Δα9) and BepA(H246A). The experiments were conducted at 30°C to slow the LptD^C^ to LptD^NC^ conversion and makes it easier to detect possible differences. Consistent with the previous results (13), BepA(H136R) had a reduced but significant ability to promote the LptD^C^ to LptD^NC^ conversion as compared with wild type BepA (Fig. 4*A* and *B*). BepA(Δα9/H136R) and BepA(H246A/H136) promoted the conversion with similar kinetics to BepA(H136R) (Fig. 4*A* and *B*). After 15 min induction, the accumulation levels of BepA and its derivatives (Fig. S5) were comparable. Also, the extracytoplasmic stress response induction by these proteins, as shown by the DegP accumulation, were similar (Fig. S5). These results indicate that the α9/H246 loop is not essential for BepA to exhibit the chaperone-like activity that promotes the LptD assembly.

**Fig. 4.**
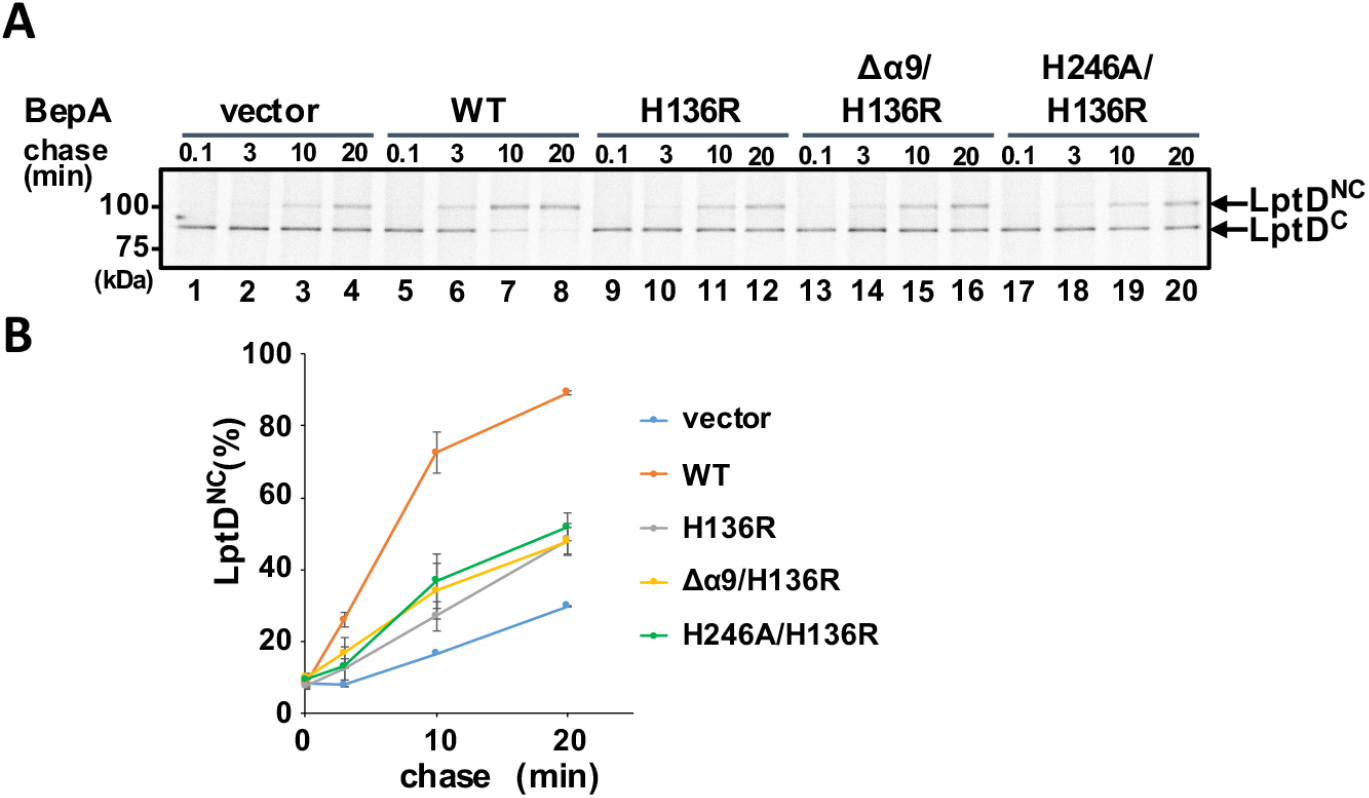
The chaperone-like activity of the α9/H246 loop mutants. (*A*) Analysis of the LptD disulfide isomerization kinetics by pulse-chase experiments. Δ*bepA* cells carrying a plasmid encoding the indicated BepA mutants were grown at 30°C in M9-based medium until early log phase, induced with 1 mM IPTG for 15 min and subjected to pulse-chase analysis as in Fig. 1*D.* The representative result of two independent replicates are shown. (*B*) LptD^NC^ (%) was calculated by dividing the band intensity for LptD^NC^ by the sum of those for LptD^C^ and LptD^NC^ and mean values are shown with standard deviations.

## Discussion

BepA is a dual functional protein that promotes assembly of LptD, but once the assembly of LptD is stagnated, it degrades the stalled/misassembled LptD (13, 14). Wild type BepA does not degrade LptD that is in the normal assembly process, indicating that the protease activity is tightly regulated so that it is expressed only as required. Recent elucidation of the full-length BepA protein structure revealed several interesting features, namely the protease active site exists inside the protease domain and is not exposed to the external milieu, and the zinc-bound water molecule is likely excluded by a His residue (His-246) that coordinates the zinc ion as the fourth ligand residue (15). His-246 is located adjacent to a short helix α9 that covers the protease active site (15). These findings suggest that some conformational change is required for BepA to express its protease activity. In this study, we focused on the α9/H246 loop, and examined its role in the BepA functions. Deletion of the α9/H246 loop or the H246A mutation increased the protease activity of BepA *in vivo* and *in vitro*. On the contrary, immobilization of the α9/H246 loop by a disulfide cross-linking repressed the proteolytic activity. These results collectively suggest that His-246 maintains BepA in the latent form and the movement of the α9/H246 loop is important for activation of BepA. Interestingly, the Δα9 and the H246A mutants of BepA degraded normally assembling LptD that is not a proteolytic substrate of wild type BepA. Thus, this auto-inhibitory mechanism should be strictly regulated to proceed appropriately in the normal functioning of BepA.

It is essential to understand how the wild type BepA in the latent (or low protease activity) form becomes activated. There are several precedents of the fourth ligand residue mediated auto-repression of the protease activity of metallopeptidases. For instances, in the latent forms of human matrix metalloproteinases (MMPs) including MMP-1 (human fibroblast collagenase) (26) and MMP-3 (human prostromelysin-I) (27), a cysteine residue in the propeptide coordinates the catalytic zinc atom instead of a water molecule. Proteolytic removal of the propeptides converts the latent proenzymes to the active forms. This auto-inhibitory Cys-mediated regulation mechanism is called a “cysteine switch”. Similar regulation mechanisms are also known for other metallopeptidases, such as fragilysin-3 of *Bacteroides fragilis* (28) and astacin of the crayfish (29). In these cases, an Asp residue in the propeptide acts as the fourth ligand, which maintains the zymogen inactive, and the enzymes are activated by removal of their propeptide. This mechanism is called an “aspartate switch”, similar to the “cysteine switch”. As for BepA, the fourth ligand that would regulate the protease activity is His-246, and thus the auto-inhibitory mechanism in this case could be called a “histidine switch”. It has been reported that an *E. coli* peptidoglycan amidase (MepA) has a His residue (His-110) that acts as the fourth metal ligand, but an Ala substitution of this residue did not elevate, but instead reduced the amidase activity of MepA (30, 31). It was thus suggested that His-110 of MepA does not act as the “histidine switch” but has a role to keep the zinc ion in the active site in the absence of a substrate. In contrast to the similarity of the auto-inhibitory mechanisms, the mechanisms of releasing the inhibition seem to be different between BepA and the cysteine/aspartate switch proteases, as the region including His-246 is not proteolytically removed from BepA during its activation. BepA not only promotes biogenesis of LptD, but also degrades a mutant form of LptD (LptD4213) that is stalled on the BAM complex and also the LptD assembly intermediate (LptD^C^) that accumulated upon depletion of LptE. Because it has been reported that LptD associates with LptE on the BAM complex in its biogenesis pathway (19, 32), the accumulated LptD^C^ molecules under the LptE-limiting condition should also be associated with the BAM complex. The protease activity of BepA should therefore be reversibly expressed to assure that BepA can degrade only the stalled or misassembled LptD on the BAM complex. It would be possible that prolonged interaction of BepA with a stalled/misassembled substrate at the BAM complex leads to transient conformational changes of the α9/H246 loop that occur stochastically or in response to some unknown signal(s) to displace His-246 from the catalytic zinc atom. His-246 in BepA is conserved among the M48 family metallopeptidases, including eukaryotic members of this family such as human OMA1 and human Ste24 protease homolog (ZMPSTE24) (Fig. S6), which raises the possibility that the residues corresponding to BepA His-246 also participate in regulation of their protease activities. In the reported structures of yeast and human Ste24 protease homologs (33, 34), the His residues corresponding to His-246 in BepA exist at similar but slightly distant locations as compared with BepA His-246 and apparently does not directly coordinate the zinc ion. The structures might represent the activated forms of these enzymes with the relocated fourth ligand His residue.

The results of our disulfide crosslinking experiments indeed demonstrate that the mobility (conformational change) of the α9/H246 region is important for the regulated expression of the BepA’s protease activity. Although it cannot be ruled out that the decreased protease activity by the cross-linking was caused by some indirect effects, such as structural distortion around the crosslinked residues, the possible flexibility of the α9/H246 region is also suggested from the structure of the *G. sulfurreducens* BepA homolog (PDB ID 3C37) in which the region around α9 is disordered. To dislocate His-246 from the active site zinc ion, it might be sufficient for the α9/H246 region to move slightly. However, in order for a substrate polypeptide to get access to the buried or recessed active site in BepA, a relatively large conformation change would be needed. The possible movement of the α9/H246 loop might be accompanied with a more dynamic conformational change to allow entry of a substrate into the active site. The structures of *E. coli* BepA and the *G. sulfurreducens* BepA homolog showed that the regions around α6, which is also not present in the minigluzincin (23), is partially disordered (Fig. S2). In addition, the self-cleavage sites (A181-L182 and A171-M172) of BepA(Δα9) and BepA(Δα9)* (*Supplementary Results*) were found to be located in the region just following α6. Taken together, these findings suggest that the region containing α6 (α6 loop) also moves flexibly to form an opening for the substrate access. Because the regions following α6 and α9 interact closely with each other in the BepA structure, the possible movement of the α6 loop and the α9/H246 loop might be coupled.

The pulse-chase experiments using the protease-dead derivatives of the Δα9 and the H246A mutants suggest that the chaperone-like activity of BepA that promotes conversion of LptD from an intermediate to the mature form can be expressed in the absence of the α9/H246 loop region. Although we cannot exclude the possibility that the α9/H246 loop is somehow involved in the chaperone-like activity, as the protease-dead mutation (H136R) partially impairs the chaperone-like activity and the α9/H246 loop might be involved in the chaperone-like activity that requires the intactness of the active site, our results indicate that the α9/H246 loop is not essential to promoting the LptD maturation. The TPR domain of BepA that has been shown to interact with LptD (14) and some regions in the BepA protease domain other than the α9/H246 loop might be implicated in the chaperone-like activity of BepA.

We propose a functional model of BepA as shown in Fig. 5. Our previous results suggest that BepA interacts with the BAM complex such that its C-terminal TPR domain is inserted into the ring-like structure of the periplasmic domain of the BAM complex (14, 15). Thus, BAM-bound BepA may accept LptD at the periplasmic side of the BAM complex, although it would be also possible that BepA first interacts with LptD in the periplasm and delivers it to the BAM complex. Therefore, BepA that interacts with LptD at the first step should be in the latent form. As mentioned above, degradation of stalled/misassembled LptD by BepA should occur on the BAM complex. Also, it is highly likely that the BepA-mediated promotion of the LptD maturation proceeds on the BAM complex; while the absence of BepA causes accumulation of the LptD assembly intermediate (LptD^C^), its conversion to the mature form (LptD^NC^) is promoted by the overexpression of LptE (13) that would form a complex with LptD on the BAM complex (19, 32), which suggest that LptD^C^ is accumulated at the BAM complex in the absence of BepA (19, 32). BepA is normally in the latent form and promotes the maturation of LptD accompanied by the disulfide rearrangement and the association with LptE on the BAM complex, resulting in release of the functional LptD/LptE complex to the OM. When the assembly of LptD is retarded on the BAM complex, the α9/H246 loop (likely in combination with the α6 loop) of BepA moves transiently in a stochastic or signal-dependent manner to activate the proteolytic activity of BepA as discussed above. After the stalled substrate is degraded and released from the BAM complex, BepA would return to the latent form. The regulated protease activity of BepA would be important even under normal growth condition as well as in the specific instance where the stalled/misassembled LptD is generated due to the *lptD* mutation or the LptE-depletion, as expression of the protease-dead BepA mutants cannot restore the drug-sensitivity of the *bepA* mutant (14). It is conceivable that BepA acts to proteolytically eliminate a small amount of stalled/misassembled LptD molecules generated under normal growth condition, which is necessary to maintain the full integrity of the OM.

**Fig. 5.**
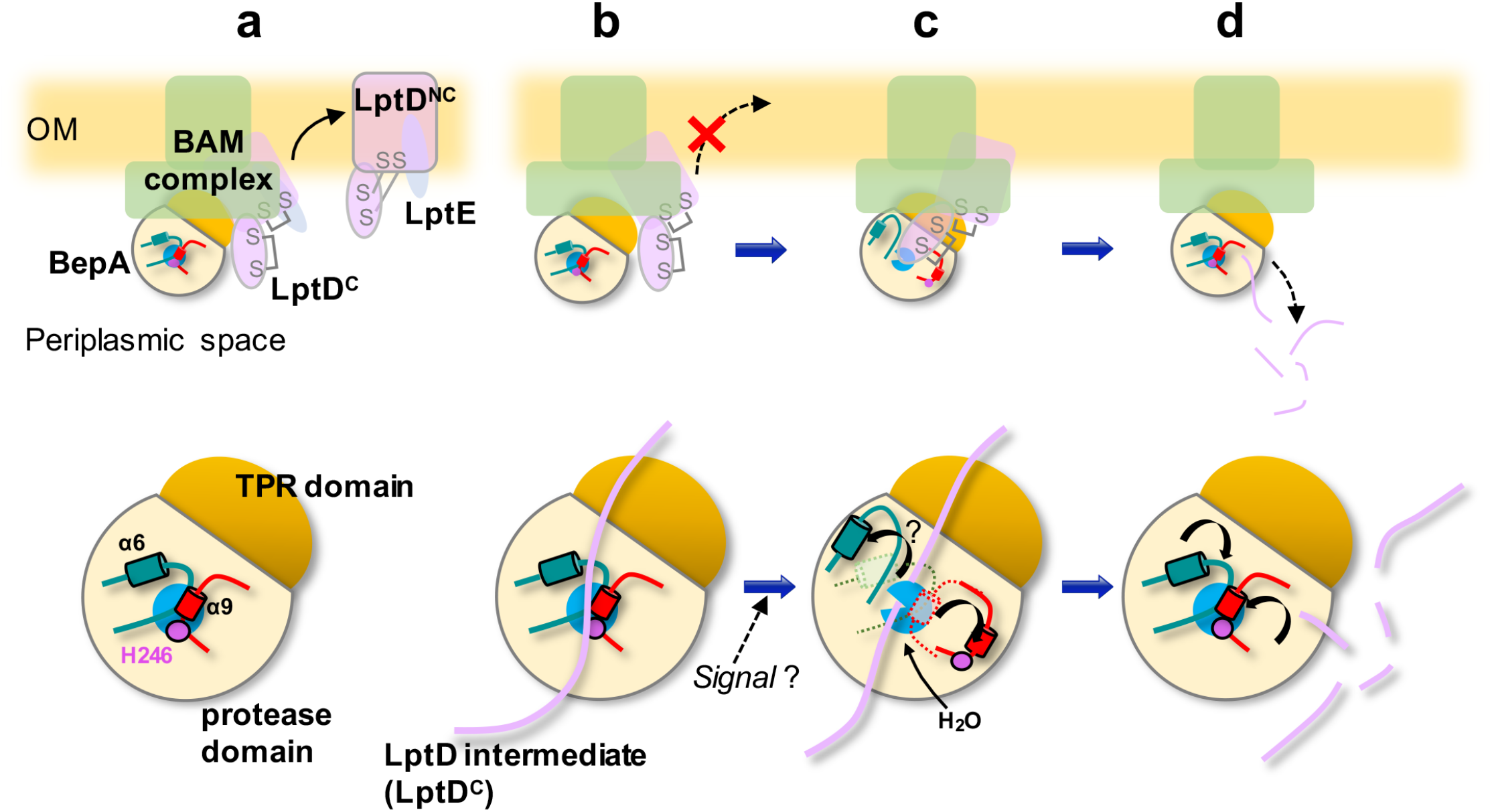
A schematic model of the reversible auto-inhibitory regulation of BepA in biogenesis and quality control of LptD. When LptD assembles normally, BepA in the latent form promotes the assembly of LptD at the BAM complex. In the latent form, the protease activity of BepA is repressed by zinc coordinating His-246 (magenta), and the access of LptD to the active site (cyan) is blocked by the α9/H246 loop (red) and the α6 loop (teal) (a). Impaired assembly of LptD causes prolonged stay of the assembly intermediate form of LptD (LptD^C^) at the BAM complex (b). Under such conditions, conformational changes of the α9/H246 loop, and possibly the α6-loop, are induced (stochastically or by some signals) to allow entry of a catalytically-required water molecule and access of the substrate to the active site (c), and the stalled LptD is degraded by BepA (d). BepA then returns to the latent form for the next round of LptD assembly/degradation (d). Note that BepA and LptD are schematically depicted as we know neither the exact regions of LptD that interacts with BepA nor the conformation of the BepA-interacting LptD. Enlarged views of BepA at each step are shown in a lower part.

We note that during the preparation of this manuscript, an archived preprint (35) reported determination of the *E. coli* BepA structure and analysis of His-246. The structure solved in that study was very similar to the structure we have reported previously and, consistent with our results, their mutational analysis of His-246 and the α9/H246 loop in that study reportedly supported the importance of these elements in the overall BepA function, although neither the protease activity nor the chaperone-like activity of BepA were directly examined in that report.

Further experiments will be needed to verify the above model. For example, the possible movability of the the α9/H246 loop and the α6 loop should be investigated experimentally by biochemical and/or biophysical approaches. Also, structural study on the BepA-substrate complex and the BepA-BAM complex as well as molecular dynamics simulation will provide useful information of the possible conformation changes of BepA. In addition, *in vitro* system to reproduce the BepA- and BAM-mediated OM assembly and degradation of LptD should be established for detailed analysis of the regulation mechanism of the BepA protease activity.

## Acknowledgements

We thank Eiji Ishii for discussion and Shin-ichi Matsuyama for antisera. This work was supported by Japan Society for the Promotion of Science KAKENHI Grants 18H023404 (to Y.A.) and 18K06136 (to S.N.), a research grant from Nagase Science and Technology Foundation (to Y.A.), and Joint Usage/Research Center program of Institute for Frontier Life and Medical Sciences, Kyoto University.

## Materials and Methods

### Bacterial strains, plasmids and media

The *E. coli K12* strains and plasmids used in this study are listed in Table S1. Details of the strain and plasmid construction and media used are described in *Supplementary Materials and Methods*.

### SDS/PAGE, Immunoblotting, Pulse-Chase and Immunoprecipitation Experiments

SDS/PAGE and immunoblotting with anti-BepA, -LptD or -DegP antiserum were carried out essentially according to the previously described procedures (36, 37). Pulse-Chase experiments with [^35^S]-Met and immunoprecipitation experiments using anti-LptD antiserum and Dynabeads Protein A (Invitrogen) were carried out essentially according to the previously described procedures (13). Details were described in *Supplementary Materials and Methods*. Anti-BepA anti serum was raised against purified BepA (13). Anti-LptD, and -DegP antisera were provided by Shin-ichi Matsuyama (Rikkyo University, Tokyo, Japan).

### Purification and In Vitro Proteolytic Activity Assay of BepA

SN896(DE3) cells carrying pCDF-bepA-his10 derivatives were grown in L medium at 30°C. BepA and its derivatives were affinity purified using a C-terminally-attached His_10_-tag essentially as described previously (13) with slight modification (see *Supplementary Materials and Methods* for details).

The reaction mixture for BepA proteolytic activity assay contained 5 mM Tris·HCl (pH 8.0), 150 mM NaCl, 1 μM ZnCl2, 7.5% glycerol, 0 or 200 μg/mL α-casein (Sigma) and 0 or 100 μg/mL BepA. Where specified, 1,10-phenanthroline or EDTA was added to the reaction mixture at the final concentration of 250 μM. After incubation at 37°C for the appropriate time periods, a portion of the samples were withdrawn, mixed with SDS sample buffer, boiled for 10 min, and subjected to 5-20% linear gradient SDS-PAGE. Proteins were visualized by staining with QC Colloidal Coomassie Stain (Bio-Rad).

## References

1. H. Nikaido, Molecular basis of bacterial outer membrane permeability revisited. Microbiol. Mol. Biol. Rev. 67, 593–656 (2003).

2. P. Sperandeo, A.M. Martorana, A. Polissi, The lipopolysaccharide transport (Lpt) machinery: A nonconventional transporter for lipopolysaccharide assembly at the outer membrane of Gram-negative bacteria. J. Biol. Chem. 292, 17981–17990 (2017).

3. T. Wu et al., Identification of a protein complex that assembles lipopolysaccharide in the outer membrane of *Escherichia coli*. Proc. Natl. Acad. Sci. U. S. A. 103, 11754–11759 (2006).

4. H. Dong et al., Structural basis for outer membrane lipopolysaccharide insertion. Nature 511, 52–56 (2014).

5. S. Qiao, Q. Luo, Y. Zhao, X.C. Zhang, Y. Huang, Structural basis for lipopolysaccharide insertion in the bacterial outer membrane. Nature 511, 108–111 (2014).

6. E. Freinkman, S.S. Chng, D. Kahne, The complex that inserts lipopolysaccharide into the bacterial outer membrane forms a two-protein plug-and-barrel. Proc. Natl. Acad. Sci. U.S.A 108, 2486–2491 (2011).

7. A.M. Plummer, K.G. Fleming, From Chaperones to the Membrane with a BAM! Trends Biochem. Sci. 41, 872–882 (2016).

8. D. Ranava, A. Caumont-Sarcos, C. Albenne, R. Ieva, Bacterial machineries for the assembly of membrane-embedded β-barrel proteins. FEMS Microbiol. Lett. 365, 1–12 (2018).

9. D.P. Ricci, T.J. Silhavy, Outer Membrane Protein Insertion by the β-barrel Assembly Machine. EcoSal Plus 8, 1–9 (2019).

10. L.R. Warner, P.Z. Gatzeva-Topalova, P.A. Doerner, A. Pardi, M.C. Sousa, Flexibility in the periplasmic domain of BamA is important for Function. Structure 25, 94–106 (2017).

11. N. Noinaj, J.C. Gumbart, S.K. Buchanan, The β-barrel assembly machinery in motion. Nat. Rev. Microbiol. 15, 197–204 (2017).

12. N.D. Rawlings et al., The MEROPS database of proteolytic enzymes, their substrates and inhibitors in 2017 and a comparison with peptidases in the PANTHER database. Nucleic Acids Res. 46, D624–D632 (2018).

13. S. Narita, C. Masui, T. Suzuki, N. Dohmae, Y. Akiyama, Protease homolog BepA (YfgC) promotes assembly and degradation of β-barrel membrane proteins in *Escherichia coli*. Proc. Natl. Acad. Sci. U. S. A. 110, E3612–E3621 (2013).

14. Y. Daimon et al., The TPR domain of BepA is required for productive interaction with substrate proteins and the β-barrel assembly machinery complex. Mol. Microbiol. 106, 760–776 (2017).

15. M. Shahrizal et al., Structural basis for the function of the β-barrel assembly-enhancing protease BepA. J. Mol. Biol. 431, 625–635 (2019).

16. G.R. Soltes, N.R. Martin, E. Park, H.A. Sutterlin, T.J. Silhavy, Distinctive roles for periplasmic proteases in the maintenance of essential outer membrane protein assembly. J. Bacteriol. 113, 8717–8722 (2016).

17. N. Ruiz, S.S. Chng, A. Hiniker, D. Kahne, T.J. Silhavy, Nonconsecutive disulfide bond formation in an essential integral outer membrane protein. Proc. Natl. Acad. Sci. U.S.A. 107, 12245–12250 (2010).

18. S.-S. Chng et al., Disulfide rearrangement triggered by translocon assembly controls lipopolysaccharide export. Science 337, 1665–1668 (2012).

19. J. Lee et al., Characterization of a stalled complex on the β-barrel assembly machine. Proc. Natl. Acad. Sci. U. S. A. 113, 8717–8722 (2016).

20. T. Hirano, N. Kinoshita, K. Morikawa, M. Yanagida, Snap helix with knob and hole: essential repeats in S. pombe nuclear protein nuc2^+^. Cell 60, 319–328 (1990).

21. L.D. D’Andrea, L. Regan, TPR proteins: the versatile helix. Trends Biochem. Sci. 28, 655–662 (2003).

22. N. Zeytuni, R. Zarivach, Structural and functional discussion of the tetra-trico-peptide repeat, a protein interaction module. Structure 20, 397–405 (2012).

23. M. Lopez-Pelegrin et al., A novel family of soluble minimal scaffolds provides structural insight into the catalytic domains of integral membrane metallopeptidases. J. Biol. Chem. 288, 21279–21294 (2013).

24. D.S. Auld, “Catalytic mechanisms for metallopeptidases” in Handbook of Proteolytic Enzymes: Third Edition, N. D. Rawlings, G. Salvesen, Eds. (Amsterdam: Elsevier Ltd, 2013), pp. 370–396.

25. T.L. Raivio, T.J. Silhavy, The σ^E^ and Cpx regulatory pathways: overlapping but distinct envelope stress responses. Curr. Opin. Microbiol. 2, 159–165 (1999).

26. D. Jozic et al., X-ray structure of human proMMP-1: New insights into procollagenase activation and collagen binding. J. Biol. Chem. 280, 9578–9585 (2005).

27. J.W. Becker et al., Stromelysin‐1: Three‐dimensional structure of the inhibited catalytic domain and of the C‐truncated proenzyme. Protein Sci. 4, 1966–1976 (1995).

28. T. Goulas, J.L. Arolas, F.X. Gomis-Rüth, Structure, function and latency regulation of a bacterial enterotoxin potentially derived from a mammalian adamalysin/ADAM xenolog. Proc. Natl. Acad. Sci. U. S. A. 108, 1856–1861 (2011).

29. T. Guevara et al., Proenzyme structure and activation of astacin metallopeptidase. J. Biol. Chem. 285, 13958–13965 (2010).

30. M. Marcyjaniak, S.G. Odintsov, I. Sabala, M. Bochtler, Peptidoglycan amidase MepA is a LAS metallopeptidase. J. Biol. Chem. 279, 43982–43989 (2004).

31. M. Firczuk, M. Bochtler, Mutational analysis of peptidoglycan amidase MepA. Biochemistry 46, 120–128 (2007).

32. G. Chimalakonda et al., Lipoprotein LptE is required for the assembly of LptD by the β-barrel assembly machine in the outer membrane of *Escherichia coli*. Proc. Natl. Acad. Sci. U. S. A. 108, 2492–2497 (2011).

33. A. Quigley et al., The structural basis of ZMPSTE24-dependent laminopathies. Science 340, 1604–1607 (2013).

34. E.E. Pryor Jr. et al., Structure of the integral membrane protein CAAX protease Ste24p. Science 339, 1600–1604 (2013).

35. J.A. Bryant et al., Structure-function analysis of the *Escherichia coli* β–barrel assembly enhancing protease BepA suggests a role for a self-inhibitory state. bioRxiv (2019) http://dx.doi.org/10.1101/689117.

36. U.K. Laemmli, Cleavage of structural proteins during the assembly of the head of bacteriophage T4. Nature 227, 680–685 (1970).

37. T. Shimoike et al., Product of a new gene, *syd*, functionally interacts with SecY when overproduced in *Escherichia coli*. J. Biol. Chem. 270 (1995).

