## Supplementary files for "Reversible auto-inhibitory regulation of *Escherichia coli* metallopeptidase BepA for selective β-barrel protein degradation"

### Supplementary Information

#### Supplementary Results

##### Isolation of the mutations that suppress self-degradation of BepA(Δα9) and BepA(H246A)

We overexpressed and purified wild-type and the mutant forms of the BepA proteins with $His_{10}$-tag at their C-terminus by metal-affinity chromatography. SDS/PAGE analysis of the eluted fraction of BepA(Δα9) and BepA(H246A) showed that the purified preparations contained less amounts of full-length BepA compared to wild-type BepA, with several faster-migrating proteins (Fig. 2*A*, lanes 5 and 11). Some of these faster-migrating species (such as 31.5 kDa and 15 kDa bands) possibly represent self-degradation products because their amounts increased upon incubation of the purified preparations with concomitant decrease in the amount of the full-length protein (Fig. 2*A*). N-terminal sequence analysis of the major fragment of about 30 kDa (Fig. 2*A*, blue arrowheads) showed that this fragment was a C-terminal part of BepA generated by cleavage between Ala-181 and Leu-182. To reduce the self-degradation, which was supposed to improve the yield of full-length BepA, we conducted random mutagenesis against the codons for A181 and L182 and screened for the mutants with decreased degradation (see *Supplementary Materials and Methods* for details). We found that a pair of mutations (A181E/L182T) significantly suppresses the *in vivo* self-degradation of BepA(Δα9) and BepA(H246A) (Fig. 2*A*, lanes 7 and 13).

#### Supplementary Materials and Methods

##### Bacterial strains, plasmids and media

Strain SN896(DE3) was constructed by lysogenizing λ(DE3) into SN896. Derivatives of pUC-bepA or pCDF-bepA-$his_{10}$ encoding a mutant form of BepA were constructed by site-directed mutagenesis using pairs of complementary primers.

Derivatives of pSTD-bepA were constructed by site-directed mutagenesis or by subcloning an EcoRI-HindIII fragment of the pUC-bepA derivatives into the same sites of pSTD689. To construct pCDF-bepA(A181E/L182T)Δ(239-247)::GSGSGS-$his_{10}$, the codons for Ala-181 and Leu-182 of the *bepA* gene on pCDF-bepAΔ(239-247)::GSGSGS-$his_{10}$ were mutagenized by site-directed mutagenesis using a pair of primers with randomized sequences for these codons. The plasmid library thus obtained was introduced into SN896(DE3) and the transformants were screened for the elevated accumulation of full-length BepA by immunoblotting. One of them that showed the highest accumulation level was stored and the DNA sequence of the *bepA* region was determined.

Unless indicated otherwise, cells were grown in L medium (10 g/L Bacto Tryptone, 5 g/L Bacto Yeast Extract and 5 g/L NaCl; pH was adjusted to 7.2 using NaOH) or M9 medium (without $CaCl_2$).

Ampicillin (50 μg/mL) and spectinomycin (each of 50 μg/mL) were added as appropriate for selecting transformants and for growing plasmid-bearing strains.

**SDS-PAGE and Immunoblotting Experiments**

Cells of SN56 and SN56/pTWV-lptD-$his_{10}$ additionally carrying a vector (pUC18 or pSTD689) or their derivatives encoding wild-type or a mutant form of BepA were grown to an early log phase at 37°C in M9 medium supplemented with 2 μg/ml thiamine, 0.2% maltose, 1 mM IPTG and either 19 amino acids other than Met or 18 amino acids other than Met and Cys. Proteins were precipitated with 5% trichloroacetic acid, washed with acetone, solubilized in SDS sample buffer with or without 2-mercaptoethanol and separated by 7.5% or 10% SDS-PAGE. Then, they were blotted onto a PVDF membrane filter (Merck Millipore; Billerica, MA, USA). The filter was blocked with 5% skimmed milk, and probed with anti-BepA, anti-LptD, or anti-DegP antiserum followed by HRP-conjugated anti-rabbit goat antibody. The proteins recognized by the antibodies were visualized using ECL Prime Western Blotting Detection Reagents (GE Healthcare) and a lumino-image analyzer (LAS-4000mini; Fujifilm).

**Pulse-Chase and Immunoprecipitation Experiments**

SN56 carrying pUC18 or its derivative encoding wild-type or a mutant form of BepA were grown at 30 or 37°C to an early log phase in M9 medium supplemented with 18 amino acids other than Met and Cys, 2 μg/ml thiamine, and 0.2% maltose. Then the cells were induced with 1 mM (final conc.) IPTG for the appropriate time periods, labeled with 370 kBq/ml [$^{35}$S]-Met (American Radiolabeled Chemicals) for 3 min at 30 or 37°C, and chased with 0.07% (final conc.) cold Met. At the indicated time points, a portion of the cultures were withdrawn and mixed with the same volume of 10% trichloroacetic acid to precipitate proteins. The proteins were dissolved in 50 μl of 50 mM Tris·HCl (pH 8.1) containing 1% SDS and 1 mM EDTA, boiled for 5 min, and diluted with 1 ml of Triton buffer containing 50 mM Tris·HCl (pH 8.1), 150 mM NaCl, 2% (wt/vol) Triton X-100, and 0.1 mM EDTA. After removal of insoluble materials by centrifugation at 20,000 × g for 5 min, the supernatant was subjected to immunoprecipitation with anti-LptD antiserum and Dynabeads Protein A (Invitrogen). Proteins recovered with the antibody were eluted from beads by boiling for 5 min in SDS sample buffer without 2-mercaptoethanol, separated by 7.5% SDS/PAGE, and visualized with with BAS-1800 (Fujifilm). Where specified, the eluted samples were further treated with 10% (final conc.) 2-mercaptoethanol before SDS/PAGE. Proteins were separated by 7.5% SDS/PAGE, and visualized with with BAS-1800 (Fujifilm). Band intensities were quantified by using MultiGauge software (Fujifilm).

**Purification of BepA.**

SN896(DE3) cells carrying pCDF-bepA-$his_{10}$ derivatives were grown in L medium at 30°C. When the culture OD (at 600 nm) reached 0.2, expression of the BepA derivatives was induced with 75 µM IPTG for 2 h. Cells were then harvested, washed once with 5 mM Tris·HCl (pH 8.0), and resuspended in 5 mM Tris·HCl (pH 8.0) containing 300 mM sucrose, 10 µg/ml DNase I and cOmplete™ EDTA-free Protease Inhibitor Cocktail (Roche). They were converted into spheroplasts by addition of 50 µg/ml lysozyme and 1 mM EDTA followed by incubation for 20 min at 4°C. After addition of 2 mM $MgCl_2$, the spheroplasts and insoluble materials were removed by successive centrifugations at 10,000 × g for 5 min and at 100,000 × g for 30 min to obtain the periplasmic fraction. The periplasmic fraction was applied to a TALON metal affinity resin (Clontech) column. The column was successively washed with buffer A [5 mM Tris·HCl (pH 8.0), 50 mM NaCl] and buffer A containing 5 mM imidazole, and finally eluted with buffer A containing 250 mM imidazole. The buffer of eluted fraction was exchanged to 5 mM Tris·HCl (pH 8.0) containing 10% glycerol by passage through a Sephadex G-25 desalting column (PD-10; GE Healthcare), followed by concentration by the Amicon® Ultra centrifugal filters (Millipore). Protein concentration of purified proteins was determined using the Bio-Rad Bradford protein assay.

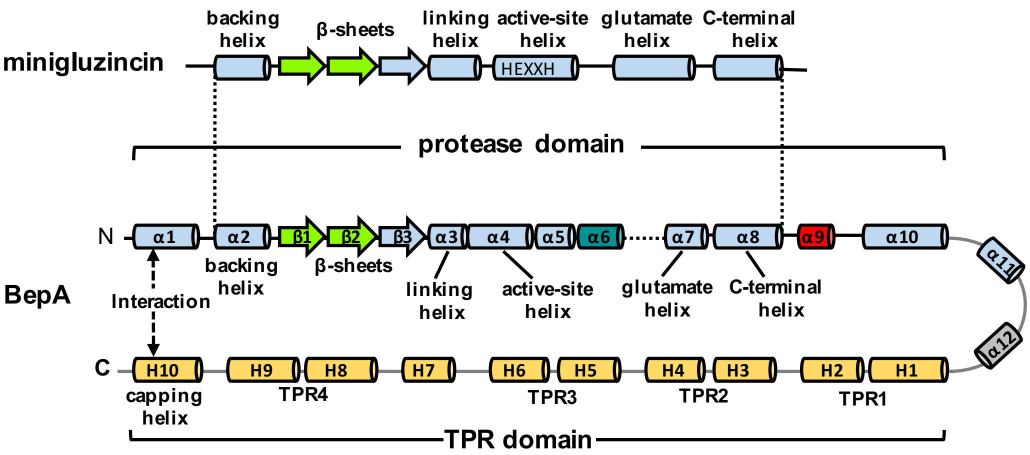


**Fig. S1.** Comparison of the secondary structure arrangements between minigluzincin and BepA. The secondary structure arrangements of the BepA (15) and minigluzincin (23) are schematically depicted. Arrows and rods indicate β-strands and α-helices, respectively, and colored as in Fig. 1*A*.

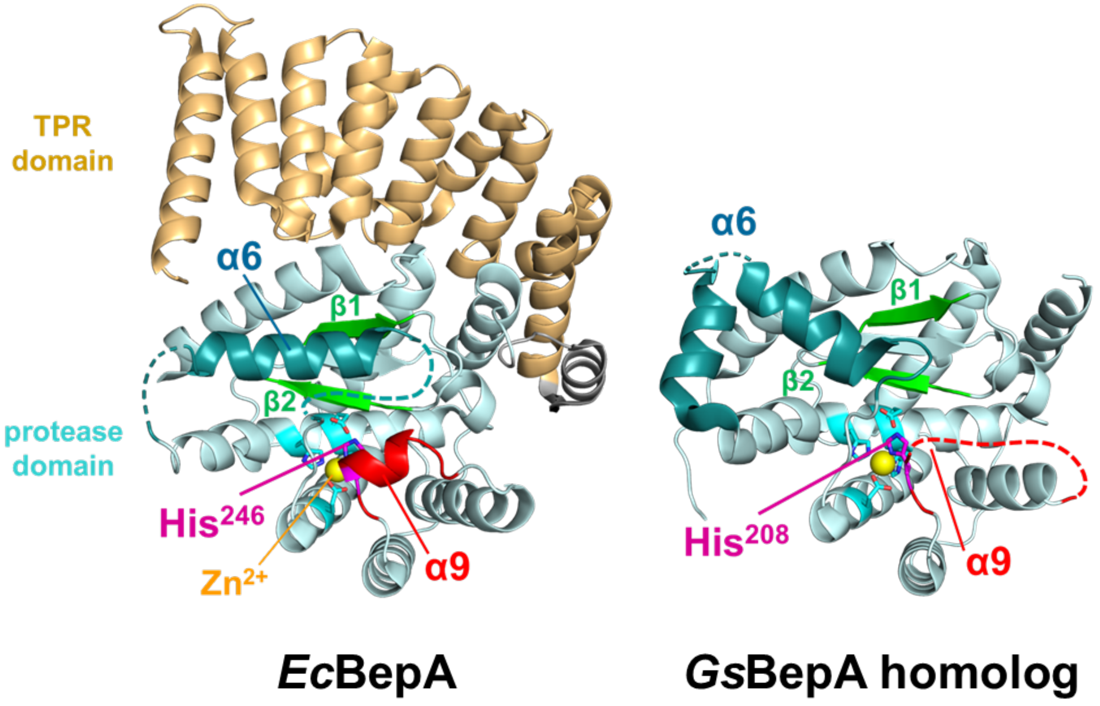


**Fig. S2.** Structures of *E. coli* BepA (PBD ID: 6AIT) and *G. sulfurreducens* BepA homolog (PDB ID 3C37). The regions corresponding to the α9/H246 loop (red), α6 loop (teal), β1/ β2 (light green), zinc atom (yellow) and His-246 residue (magenta) of *E. coli* BepA are colored as in Fig. 1*A*.

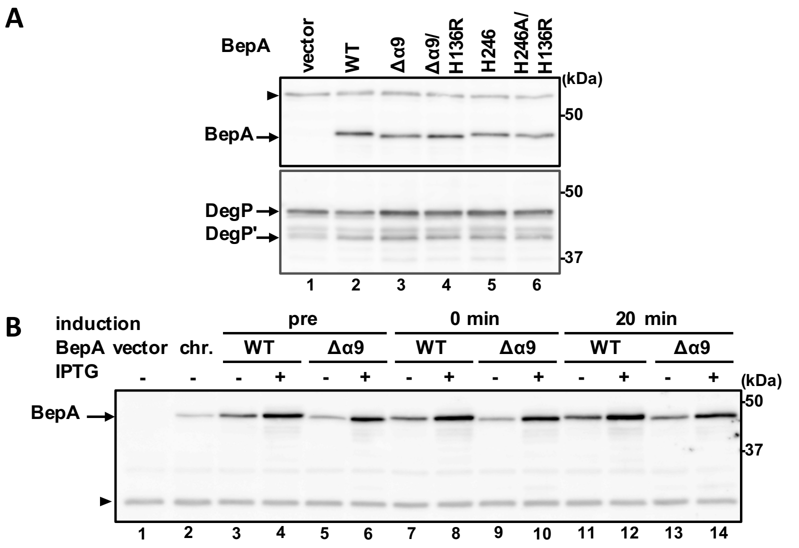


**Fig. S3.** Accumulation levels of BepA and DegP under the conditions of the pulse-chase experiments shown in Fig. 1. (*A*) Cells were grown and treated as in Fig. 1*D* except that they were not pulse-labeled. A portion was withdrawn at the time point corresponding to just before pulse-labeling or at 60 min chase for analysis of BepA and DegP, respectively. Proteins were acid-precipitated and subjected to SDS/PAGE and immunoblotting analysis with anti-BepA (upper panel) or anti-DegP (lower panel) antiserum. DegP' indicates the degradation products of DegP. The migration positions of molecular mass markers are shown (*B*) Cells were grown and treated as in Fig. 1*E* except that they were not pulse-labeled. A portion was withdrawn at the time point corresponding to just before (IPTG -) or 15 minutes after (IPTG +) the induction for each culture. Proteins were acid-precipitated and subjected to SDS/PAGE and immunoblotting analysis with anti-BepA antiserum. Arrowheads indicate a non-specific band serving as a loading control. The representative results of two independent replicates are shown.

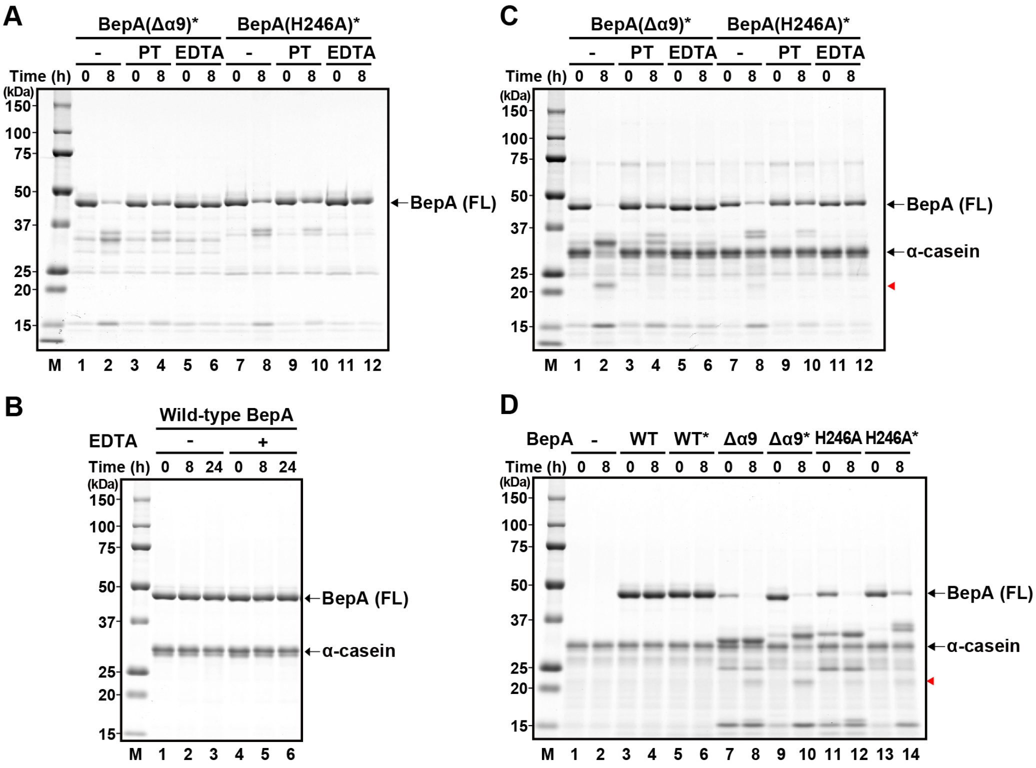


**Fig. S4.** Effects of metal chelators on the proteolytic activity of the Δα9 and the H246A mutants. (*A*) Effects of metal chelators on the self-cleavage activity of BepA mutants were analyzed by incubating the BepA(Δα9)* or the BepA(H246A)* protein, each carrying the A181E/L182T mutations, at 37°C in the absence (-) or presence of 250 μM 1,10-phenanthroline (PT) or 250 μM EDTA for 0 or 8 h, followed by SDS/PAGE and CBB staining. (*B*) Wild-type BepA with a C-terminus $His_{10}$-tag was incubated with α-casein at 37°C in the absence (-) or presence (+) of 250 μM EDTA for 0, 8 or 24 h and analyzed by SDS/PAGE and CBB staining. (*C*) Effects of metal chelators on the caseinolytic activity of BepA(Δα9)* or BepA(H246A)* were analyzed as in *B* except that the reaction mixture contained α-casein. (*D*) Effects of the A181E/L182T mutations on the caseinolytic activity of BepA were analyzed by incubating wild-type (WT) or the mutant forms of the BepA proteins, either with or without the A181E/L182T mutation, with α-casein at 37°C for 0 or 8 h followed by SDS/PAGE and CBB staining. Red arrowheads indicate a degradation product of α-casein.

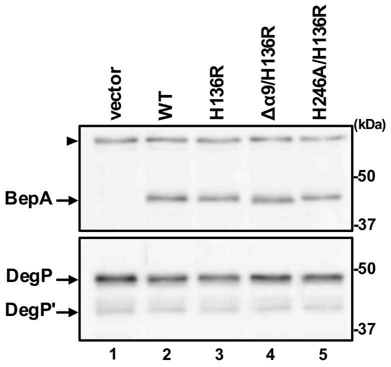


**Fig. S5.** Accumulation levels of BepA and DegP under the conditions of the pulse-chase experiments shown in Fig. 4. Cells were grown and treated as in Fig. 4 except that they were not pulse-labeled. A portion was withdrawn at the time point corresponding to just before pulse-labeling or at 20 min chase time for analysis of BepA (upper panel) and DegP (lower panel), respectively. Proteins were acid-precipitated and subjected to SDS/PAGE and immunoblotting analysis with anti-BepA or anti-DegP antiserum. Arrowhead indicates non-specific bands serving as a loading control. The representative results of two independent replicates are shown.

```
                                  HExxH motif
Ec BepA    116  LHSALFRYSDNESQLASVMAHEISHVTQRHLARAMEDQQRSAPLTWVGALGSILLAM----ASPQAGMAALT  183
Ec HtpX    120  VSTGLLQNMSPD-EAEAVIAHEISHIANGDMVTMTLIQGVVNTFV---IFISRIL-A----QLAAGFMGGNR  182
Ec LoiP    111  VYSGLMDMMTDN-EVEAVIGHEMGHVALGHVKKGMQVA--LGTNA------VRV-AA----ASAGGIVGSLS  168
Ec YcaL    115  VYSGLMDMMNDN-EIEGVLGHELGHVALGHSLAEMKAS--YAIVA------ARD-AI----SATSGVASQLS  172
Hs OMA1    307  VFTGFLNSVTDIHQLSFLLGHEIAHAVLGHAAEKAGMVHLLDFLG---MIFLTMIWA----ICPRDSLALLC  371
Hs STE24   315  AKVKNKKQGCKNEEVLAVLGHELGHWKLGHTVKNIIISQMNSFLCF---FLFAVLIGRKELFAAFGFYDS--  381
                             :   ::.**:.*    .
```

```
                                                    Third ligand
Ec BepA    184  ---------------------GTLAGTRQ--GM--ISFTQQNEQEADRIGIQVLQRSGFDPQAMPTFLEKL-  229
Ec HtpX    183  DEGEESNGNPLIYFAVATVLELVF-GILA--SIITMWFSRHREFHADAGSAKLVGREKM-IAALQRLKTSYE  250
Ec LoiP    169  --------------------QSQLGNLGE--KLVNSQFSQRQEAEADDYSYDLLRQRGISPAGLA---TSF-  214
Ec YcaL    173  --------------------RSQLGDIAE--GAINAKYSRDKESEADDFSFDLLKKRGISTQGLV---GSF-  218
Hs OMA1    372  --------------------QWIQSKLQE--YMFNRPYSRKLEAEADKIGLLLAAKACADIRASSVFWQQM-  420
Hs STE24   382  --QPTLIG-------LLIIFQFIFSPYNEVLSFCLTVLSRRFEFQADAFAKKLGKAKDLYSA-LIKLNK---  440
                                                      ::  * .**  .  :
```

```
                                         His246
Ec BepA    230  ---------LDQARYSSRPPEILLTHPLPESRLADARNRANQMRPMVVQSSEDFYLAKARTLGMYN  286
Ec HtpX    251  PQEATSMMALCINGKSKSLSELFMTHPPLDKRIEALRTGEYLK-----------------------  293
Ec LoiP    215  -------EKLAKLEEGRQS-SMFDDHPASAERAQHIRDRMSADGIK--------------------  252
Ec YcaL    219  -------ETLASLDGGRTQ-SMFDSHPPSTERAQHIRDRIASGK----------------------  254
Hs OMA1    421  -------EFVDSLHGQPKMPEWLSTHPSHGNRVE-YLDRLIPQALKIREMCNCPPLSNPDPRLLFK  478
Hs STE24   441  -------DNLGFPVSDWLFSMWHYSHPPLLERLQALKTMKQH------------------------  475
                         :               **   .*
```

**Fig. S6.** Sequence alignment of M48 peptidase family proteins. Amino acid sequences of *Escherichia coli* BepA (Ec BepA; UniProt P66948), *E. coli* HtpX (Ec HtpX; UniProt P23894), *E. coli* LoiP (Ec LoiP; UniProt P25894), *E. coli* YcaL (Ec YcaL; UniProt P43674) and *Homo sapiens* OMA1 (Hs OMA1; UniProt Q96E52) and *H. sapiens* ZMPSTE24 (Hs STE24; UniProt O75844) are aligned by the Clustal Omega program (https://www.ebi.ac.uk/Tools/msa/clustalo/). Conserved residues are colored in red.

### Table S1: Strains and plasmids used in this study

| ***E. coli* strains** | **Genotype** | **Reference** |
|---|---|---|
| AD16 | Δ*pro-lac thi*/ F'$lacI^q$ ZΔM15 $Y^+$ $pro^+$ | (1) |
| SN56 | AD16 Δ*bepA*::FRT | (2) |
| MC4100 | *araD139* Δ(*argF-lac*)*U169 rpsL150 relA1 flbB5301 deoC1 ptsF25 rbsR* | (3) |
| SN896 | MC4100 Δ*bepA*::FRT | (2) |
| SN896(DE3) | SN896, λDE3 | (2) |
| **Plasmids** | **Description** | **Reference or source** |
| pUC18 | Expression vector; $P_{lac}$, *bla* | (4) |
| pUC-bepA | pUC18 derivative encoding BepA | (2) |
| pUC-bepA(H136R) | pUC-bepA derivative, H136R | (2) |
| pUC-bepA(E137Q) | pUC-bepA derivative, E137Q | (2) |
| pUC-bepAΔ(239-247)::GSGSGS | pUC-bepA derivative, Δα9 | This study |
| pUC-bepA(H136R)Δ(239-247)::GSGSGS | pUC-bepA derivative, H136R, Δα9 | This study |
| pUC-bepA(E137Q)Δ(239-247)::GSGSGS | pUC-bepA derivative, E137Q, Δα9 | This study |
| pUC-bepA(H246A) | pUC-bepA derivative, H246A | This study |
| pUC-bepA(H136R/H246A) | pUC-bepA derivative, H136R/H246A | This study |
| pUC-bepA(E137Q/H246A) | pUC-bepA derivative, E137Q/H246A | This study |
| pSTD689 | Expression vector; Plac, *aadA* | (5) |
| pSTD-bepA | pSTD689 derivative encoding BepA | (6) |
| pSTD-bepA(H136R) | pSTD-bepA derivative, H136R | This study |
| pSTD-bepAΔ(239-247)::GSGSGS | pSTD-bepA derivative, Δα9 | This study |
| pSTD-bepA（H136R）Δ(239-247)::GSGSGS | pSTD-bepA derivative, H136R, Δα9 | This study |
| pSTD-bepA(H246A) | pSTD-bepA derivative, H246A | This study |
| pSTD-bepA(H136R/H246A) | pSTD-bepA derivative, H136R/H246A | This study |
| pSTD-bepA(E103C) | pSTD-bepA derivative, E103C | This study |
| pSTD-bepA(E241C) | pSTD-bepA derivative, E241C | This study |
| pSTD-bepA(E103C/E241C) | pSTD-bepA derivative, E103C/E241C | This study |
| pTWV228 | Expression vector; Plac, *bla* | (7) |
| pTWV-lptD-his$_{10}$ | pTWV228 derivative encoding LptD-His$_{10}$ | (6) |
| pCDFDuet-1 | Expression vector; PT7 *aadA* | Novagen |
| pCDF-bepA-his$_{10}$ | pCDFDuet-1 derivative encoding BepA-His$_{10}$ | (2) |
| pCDF-bepA(A181E/L182T)-his$_{10}$ | pCDFD-bepA-his$_{10}$ derivative, A181E/L182T | This study |
| pCDF-bepAΔ(239-247)::GSGSGS-his$_{10}$ | pCDFD-bepA-his$_{10}$ derivative, Δα9 | This study |
| pCDF-bepA(A181E/L182T)Δ(239-247)::GSGSGS-his$_{10}$ | pCDFD-bepA-his$_{10}$ derivative, A181E/L182T, Δα9 | This study |
| pCDF-bepA( H136R)(A181E/L182T)Δ(239-247)::GSGSGS-his$_{10}$ | pCDFD-bepA-his$_{10}$ derivative, H136R/A181E/L182T, Δα9 | This study |
| pCDF-bepA(H246A)-his$_{10}$ | pCDFD-bepA-his$_{10}$ derivative, H246A | This study |
| pCDF-bepA(A181E/L182T)(H246A)-his$_{10}$ | pCDFD-bepA-his$_{10}$ derivative, A181E/L182T/H246A | This study |
| pCDF-bepA(H136R)(A181E/L182T)(H246A)-his$_{10}$ | pCDFD-bepA-his$_{10}$ derivative, H136R/A181E/L182T/H246A | This study |


1. A. Kihara, Y. Akiyama, K. Ito, A protease complex in the *Escherichia coli* plasma membrane: HflKC (HflA) forms a complex with FtsH (HflB), regulating its proteolytic activity against SecY. *EMBO J.* **15**, 6122–6131 (1996).
2. S. Narita, C. Masui, T. Suzuki, N. Dohmae, Y. Akiyama, Protease homolog BepA (YfgC) promotes assembly and degradation of β-barrel membrane proteins in *Escherichia coli*. *Proc. Natl. Acad. Sci. U.S.A.* **110**, E3612–E3621 (2013).
3. T.J. Silhavy, M.L. Berman, L.W. Enquist, Experiments with gene fusions. *Cold Spring Harb. Lab. New York* (1984).
4. C. Yanisch-Perron, J. Vieira, J. Messing, Improved M13 phage cloning vectors and host strains: nucleotide sequences of the M13mp18 and pUC19 vectors. *Gene* **33**, 103–119 (1985).
5. K. Kanehara, K. Ito, Y. Akiyama, YaeL proteolysis of RseA is controlled by the PDZ domain of YaeL and a Gln-rich region of RseA. *EMBO J.* **22**, 6389–6398 (2003).
6. Y. Daimon *et al.*, The TPR domain of BepA is required for productive interaction with substrate proteins and the β-barrel assembly machinery complex. *Mol. Microbiol.* **106**, 760–776 (2017).
7. T. Taura, Y. Akiyama, K. Ito, Genetic analysis of SecY: Additional export-defective mutations and factors affecting their phenotypes. *Mol. Gen. Genet.* **243**, 261–269 (1994).
